# Explanation implies causation?

**DOI:** 10.1101/218784

**Authors:** Leslie Myint, Jeffrey T. Leek, Leah R. Jager

## Abstract

Most researchers do not deliberately claim causal results in an observational study. But do we lead our readers to draw a causal conclusion unintentionally by explaining why significant correlations and relationships may exist? Here we perform a randomized study in a data analysis massive online open course to test the hypothesis that explaining an analysis will lead readers to interpret an inferential analysis as causal. We show that adding an explanation to the description of an inferential analysis leads to a 15.2% increase in readers interpreting the analysis as causal (95% CI 12.8% - 17.5%). We then replicate this finding in a second large scale massive online open course. Nearly every scientific study, regardless of the study design, includes explanation for observed effects. Our results suggest that these explanations may be misleading to the audience of these data analyses.

## Main Text

Facebook causes cancer(*1*), drinking too much tea causes prostate cancer (*2*), eating chocolate helps people stay thin(*3*). We all know that correlation doesn’t imply causation, but we’ve also all seen exaggerated headlines in the media that don’t quite capture the true results of a scientific study. A recent report in the British Medical Journal found the fault may not lie entirely with the media(*4*), but may be aided by exaggerated press releases from universities themselves. In fact, in their study of 462 press releases, the study authors found that 33% (26% to 40%) contained exaggerated causal claims. Regardless, of where the exaggeration happens, a result seems more realistic if you can explain why you think it is happening.

Most researchers do not deliberately claim causal results in an observational study. But do we lead our readers to draw a causal conclusion unintentionally by explaining why significant correlations and relationships may exist? Once we discover that an association exists, it’s natural to want to explain why it does. We may describe potential mechanisms, make connections to previous literature, or put an observation in context. Despite these explanations, causal relationships are not proven in a single observational study and are only increasingly substantiated over the course of many such studies. There is observational evidence suggesting a noticeable prevalence of inappropriate causal language in both nutritional (*5*) and educational (*6*) research studies. Here we report the results of a randomized experiment performed on an online educational platform that suggest a strong effect of explanatory language on students’ perception of whether a study is correlation or causation.

Different types of studies have different analysis goals (Table 1) (*7*). We were interested in whether people can distinguish between a study whose goal was inferential and one whose goal was actually causal, as this is a common error often termed “correlation does not equal causation”. We wanted to know whether including language explaining an observed association, leads people to believe that an inferential study is causal. To test this hypothesis, we ran an experiment in a large online open-access data analysis course. This course was an introductory-level course that covered basic data analytic concepts. Our experiment involved a single randomized quiz question administered during the course. We originally ran the experiment in January 2013, but later independently replicated our experiment in a separate offering of the course in October 2013. Between these two replications, over 22,000 students completed versions of our experimental question.

**Table 1:** Goals for different analysis types (7).

| Type of analysis | Goal of analysis |
| --- | --- |
| Descriptive | Summarizing the data without interpretation |
| Exploratory | Summarizing the data with interpretation, but without generalization beyond the original sample |
| Inferential | Generalizing beyond the original sample, with the goal of describing an association in a larger population |
| Predictive | Generalizing beyond the original sample, with the goal of predicting a measurement for a new individual |
| Causal | Generalizing beyond the original sample, with the goal of learning how changing the average of one measurement affects, on average, another measurement |
| Mechanistic | Generalizing beyond the original sample, with the goal of learning how changing one measurement deterministically affects another variable's measurement. |

Early in the course, students were presented with the definitions of six possible types of data analysis (descriptive, exploratory, inferential, predictive, causal, and mechanistic) consistent with those shown in Table 1. In the subsequent course quiz, we provided students with an description of an inferential study - from which we can only infer correlation:

> *We take a random sample of individuals in a population and identify whether they smoke and if they have cancer. We observe that there is a strong relationship between whether a person in the sample smoked or not and whether they have lung cancer. We claim that the smoking is related to lung cancer in the larger population.*

We randomized students to see or not see an explanatory interpretation accompanying this description. Students in this explanatory interpretation group saw an additional sentence:

> *We explain we think that the reason for this relationship is because cigarette smoke contains known carcinogens such as arsenic and benzene, which make cells in the lungs become cancerous.*

All students were then asked to identify the type of analysis for these results. In addition to the correct answer (inferential), students were presented at random with three of four possible incorrect answer choices (descriptive, causal, predictive, mechanistic). That is, approximately 25% of students made their choice from inferential, descriptive, causal, and predictive, approximately 25% from inferential, descriptive, causal, and mechanistic, and so on. Although the described analysis is inferential in nature, we hypothesized that students who saw the explanatory language would be more likely to identify the analysis as causal if given that choice. Because students were able to retake this quiz multiple times in order to achieve a passing grade, we collected answers from each student’s first attempt (Table 2).

In our original experiment (January 2013), 20,256 students completed our experimental quiz question. We present the results for two groups of students: (1) those who chose between inferential, causal, predictive, and mechanistic analyses and (2) those weren’t given causal as a choice, but instead chose between inferential, descriptive, predictive, and mechanistic analyses. The results for the other student groups can be found in the Supplementary Information.

Among students selecting from inferential, causal, predictive, and mechanistic answer choices, the majority (68.5%) correctly answered that the description referred to an inferential data analysis (Table 2). However, a significantly higher percentage of students who were shown the explanatory language claimed it was a causal analysis compared to students who did not see the additional language: 31.8% compared to 16.6% (95% CI for difference: 12.8% - 17.5%). These results indicate that explanatory language increases the chance a student will mistake an inferential result as causal. In this case students who saw the additional explanation were almost twice as likely to claim the results as causal.

**Table 2:**
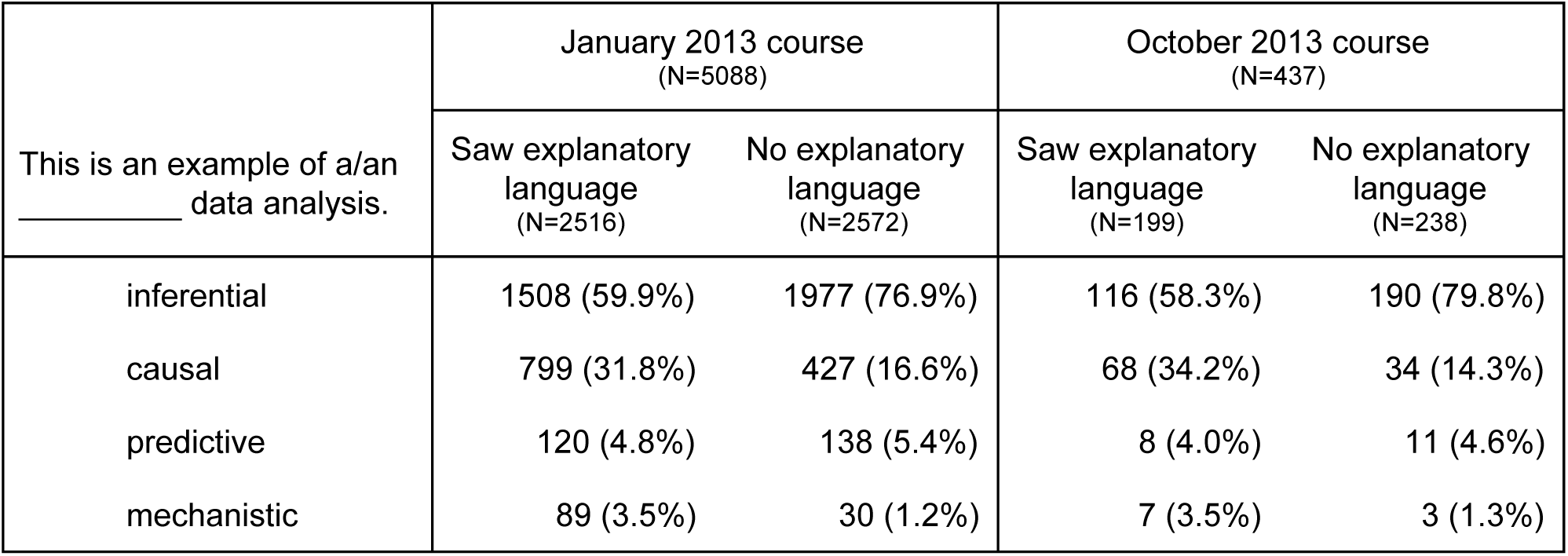
Student results for randomized quiz question asking them to identify the type of data analysis. The quiz question described an inferential analysis. Students were randomized to see or not see explanatory language that hypothesized why the association occurred.

| This is an example of a/an _____ data analysis. | January 2013 course<br>(N=5088) |  | October 2013 course<br>(N=437) |  |
| --- | --- | --- | --- | --- |
|  | Saw explanatory language<br>(N=2516) | No explanatory language<br>(N=2572) | Saw explanatory language<br>(N=199) | No explanatory language<br>(N=238) |
| inferential | 1508 (59.9%) | 1977 (76.9%) | 116 (58.3%) | 190 (79.8%) |
| causal | 799 (31.8%) | 427 (16.6%) | 68 (34.2%) | 34 (14.3%) |
| predictive | 120 (4.8%) | 138 (5.4%) | 8 (4.0%) | 11 (4.6%) |
| mechanistic | 89 (3.5%) | 30 (1.2%) | 7 (3.5%) | 3 (1.3%) |

This increase in the choice of a causal analysis when faced with explanatory language corresponded to a decrease in choice of an inferential analysis. The percentages of students who chose either a predictive or descriptive analysis was similar between the two treatment groups. However, there was an increase in the percentage of students who claimed the result was mechanistic in the explanatory language group: 3.5% compared to 1.2%. This isn’t unexpected, since a mechanistic result is similar to a causal result in that is describes a deterministic process by which one variable affects another.

Among students who weren’t given the option to select a causal as an answer (selecting instead from inferential, predictive, descriptive, and mechanistic analyses), a higher percentage (84.6%) correctly answered that the description referred to an inferential data analysis (Table 3). In this case, a significantly higher percentage of students correctly claimed the analysis was inferential when not shown the explanatory language: 88.2% compared to 80.9% (95% confidence for the difference: 5.2% - 9.3%). These results indicate that, even without the ability to identify the analysis as causal, students had a harder time correctly identifying an inferential study when given hypothesized information about the reason for a correlation. The size of the effect is must smaller than with the causal answer option, however. The decrease in correct answers again corresponded to an increase in choice of a mechanistic analysis.

**Table 3:**
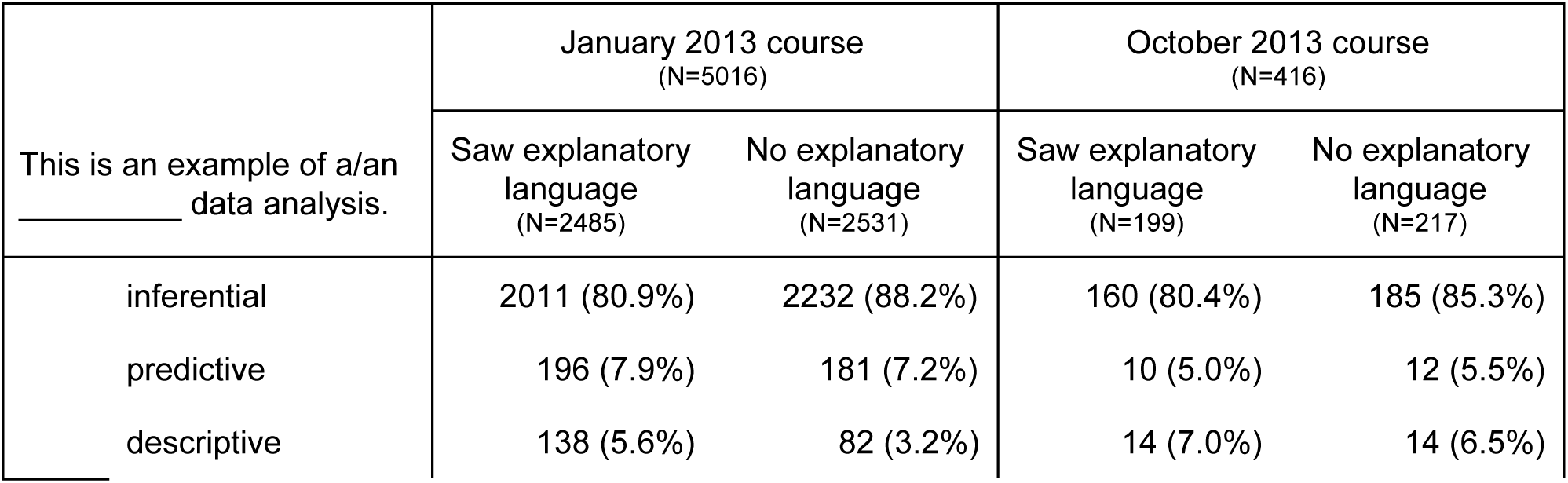
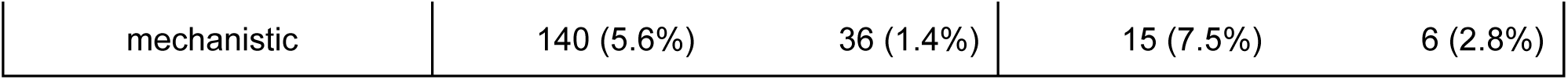
Student results when “causal” was not an answer choice.

| This is an example of a/an _____ data analysis. | January 2013 course<br>(N=5016) |  | October 2013 course<br>(N=416) |  |
| --- | --- | --- | --- | --- |
|  | Saw explanatory language<br>(N=2485) | No explanatory language<br>(N=2531) | Saw explanatory language<br>(N=199) | No explanatory language<br>(N=217) |
| inferential | 2011 (80.9%) | 2232 (88.2%) | 160 (80.4%) | 185 (85.3%) |
| predictive | 196 (7.9%) | 181 (7.2%) | 10 (5.0%) | 12 (5.5%) |
| descriptive | 138 (5.6%) | 82 (3.2%) | 14 (7.0%) | 14 (6.5%) |
| mechanistic | 140 (5.6%) | 36 (1.4%) | 15 (7.5%) | 6 (2.8%) |

To confirm our results, we performed an independent replication of our experiment in a later offering of the same data analysis course. In the replication (October 2013), 1762 students completed our experimental quiz question. The results of this replication were consistent with those in the original experiment (Table 2 and 3 and Supplementary Information). For students with inferential, causal, predictive, and mechanistic answer choices, the percentage of those who claimed it was a causal analysis was significantly higher among those who saw the explanatory language: 34.2% compared to 14.3% (95% CI for difference: 11.4% - 28.3%). For students without a causal answer choice, a higher percentage correctly identified the analysis as inferential when not shown the additional explanatory language, but the result was not statistically significant: 85.3% compared to 80.4% (95% CI for difference: -2.9% - 12.6%). The results of this replication show nearly the same effect of explanatory language on the chance that a student will interpret an inferential analysis as causal.

We know that the way data is visualized can affect how well people derive information from graphs (*8*). The results of this experiment suggest that the way we write about a data analysis is also critical. We have shown a clear causal effect of explanatory statements on perceptions of research results and replicated the effect in a second experiment. In both academic and mainstream scientific writing, there is a desire to put results into context with hypothesized mechanistic explanations to enhance the narrative around a set of empirical results. Nearly every study includes this type of explanation in the discussion section. However, our results suggest that such efforts may actually cause readers to be misled about the strength of the scientific evidence. The misinterpretation may be exacerbated by the phenomenon that readers are swayed to believe a statement when they are told scientists understand it (*9*). It turns out when talking about empirical evidence we collect through simple observation, it may be dangerous to explain.

The code and data used to perform this analysis are available at:

https://github.com/leekgroup/explanatory_language

